# Impaired Sertoli-Spermatogonia Interactions Contribute to Oligospermia and Infertility in F1 Captive-bred Male *Solea senegalensis*

**DOI:** 10.1101/2025.07.17.664418

**Authors:** Guillermo Barturen, Diego Robledo, Francisca Robles, Rose Daniels, Maialen Caballeda, Dorinda Torres-Sabino, Rafael Navajas-Pérez, Paulino Martínez, Carmelo Ruiz-Rejón, Roberto De la Herrán

## Abstract

Reproductive dysfunction of captive-bred males of the Senegalese sole (*Solea senegalensis*) represents a significant bottleneck for its aquaculture, as these fish exhibit reduced sperm production and impaired fertility compared to wild-bred counterparts acclimated to farm conditions. To elucidate the cellular and molecular mechanisms underlying this phenomenon, single-nuclei RNA sequencing was performed on gonadal tissue from adult captive-bred and wild-bred males. The analysis yielded a high-quality dataset comprising ∼80.000 cells, which were grouped into eleven distinct clusters representing all major germline and somatic cell types, including spermatogonial stem cells, differentiating spermatogonia, spermatocytes, spermatids, Sertoli cells, Leydig cells, immune cells, and peritubular myoid cells. It is noteworthy that captive-bred males exhibited a marked overrepresentation of proliferative spermatogonia and a significant reduction in mature spermatids, suggesting a disruption in the progression of spermatogenesis. Differential expression and functional enrichment analyses revealed that spermatogonial cells in captive-bred males displayed heightened translational activity alongside downregulation of pathways related to cell-cell communication and interaction. Focused cell-cell communication analyses further indicated defective Sertoli-spermatogonia interactions as a key factor contributing to oligospermia and infertility of captive-bred males. This study provides the first single-nuclei transcriptomic atlas of the Senegalese sole male gonad, offering valuable insights into the molecular basis of reproductive failure in captivity related to gonadal development. The findings of the study will inform future strategies to enhance selective breeding and improve aquaculture productivity for this economically important species.

## Introduction

Adaptation to the captive aquaculture environment entails complex changes in animal behavior, physiology, and welfare, requiring species-specific strategies to ensure optimal acclimation^1,2^. Key challenges in aquaculture production include nutrition, disease management, and, critically, reproductive success^3^. Therefore, establishing an efficient captive breeding system is a fundamental prerequisite for sustainable production. This requires not only the optimization of management protocols, such as spawning, incubation and hatchery practices, but also consideration of both intentional and unintentional selection pressures associated with domestication^4,5^. Nevertheless, reproductive dysfunctions are frequently observed in captive fish populations, including delayed or incomplete gametogenesis, reduced levels of sex hormones, smaller and lighter eggs, and diminished reproductive success, often associated to abnormal behaviors^6^. Effective breeding programs, which are essential in the competitive aquaculture industry, can only be implemented once these reproductive challenges are successfully addressed^7^.

Flatfish aquaculture includes several commercial valuable species, such as turbot (*Scophthalmus maximus*), Japanese flounder (*Paralichthys olivaceus*), and tongue sole (*Cynoglossus semilaevis*), for which selective breeding programs supported by standardized reproductive protocols have been successfully established^8,9^. Among European aquaculture candidates, the Senegalese sole (*Solea senegalensis*) stands out as one of the most promising species due to its excellent flesh quality and high market value^10^. However, reproduction remains a major challenge, as captive-bred males exhibit reproductive dysfunction, most notably the failure to perform courtship behavior. In addition, males that have been born and raised in captivity, captive-bred animals or usually called F1^11^ (CB), have been reported to produce low and variable sperm volumes^12–14^, significantly lower than their wild counterparts^13^, which hampers the use of in vitro fertilization techniques and limits the implementation of efficient selective breeding programs. In contrast, wild-bred (WB) males acclimated to captive conditions can reproduce successfully^11,15^. Notably, this reproductive dysfunction is not observed in CB females, which are able to reproduce with WB males^15^. This sex-specific impairment in CB males presents a significant challenge to the sustainability and competitiveness of the Senegalese sole aquaculture industry.

To investigate the causes of reproductive failure in CB Senegalese sole males, various approaches have been undertaken, including studies on molecular clocks involved in feeding, photoperiod regulation, and reproduction^16–18^, as well as investigations into the endocrine mechanisms controlling gonadal development and maturation^19–21^. However, the molecular mechanism underlying the abnormal gonadal maturation and reproduction failure in CB males remain poorly understood. Currently, genomic resources for the Senegalese sole are now well developed, including a highly contiguous, chromosome-level genome assembly with comprehensive annotation, providing a solid foundation for investigating the genetic and regulatory factors underlying reproduction^22–24^. In addition, recent advances in high-throughput genomics, particularly single-nuclei transcriptomics, offer powerful tools to unravel the cellular and molecular basis of these dysfunctions by enabling detailed characterization of gonadal differentiation and gene expression at single-nuclei resolution.

In this study, we conducted a single-nuclei RNA sequencing (snRNA-Seq) of the gonads from adult *S. senegalensis* males, comparing two CB and two WB individuals. Sexually mature fish were chosen to enable a direct comparison of gonadal cell composition and gene expression profiles between WB males, capable of successful reproduction in captivity, and CB males, which exhibit reduced reproductive performance. To our knowledge, this represents the first single-nucleil transcriptomic analysis of the male gonad in *S. senegalensis*. This study lays the groundwork for future research aimed at resolving reproductive dysfunction and enhancing the aquaculture production of this species, highlighting the potential of single-nuclei genomics to inform targeted interventions and improve selective breeding outcomes.

## Results

### Single-nuclei RNA-Seq analysis of *S. senegalensis* gonads yielded a good number of high-quality cells

Single-nuclei RNA-Seq analysis using the 10X Genomics protocol was performed on gonads from two CB and two WB acclimated to farm conditions *S. senegalensis* males. After background removal with Cell Ranger^25^ and filtering out cells with fewer than 500 reads, 46,757 and 38,749 cells from CB and WB animals, respectively, were included in the analysis. Additionally, 3,476 genes with more than 3 reads in at least 50 cells were selected for further analyses.

Only a few cells showed an elevated percentage of mitochondrial RNA, which indicates a low percentage of dying cells (Supplementary Figure 1A). After filtering out cells with mitochondrial RNA content above 10% and ribosomal RNA content below 1%, few cells were removed, barely noticeable in the quality control plots after removing these low-quality cells (Supplementary Figure 1B). For three of the samples (CB2, WB1 and WB2) sequencing saturation in terms of number of unique genes per sequenced molecules was reached, and most of the cells aligned to the best fit line showing a good relationship between sequenced depth and number of genes as expected in good quality cells (Supplementary Figure 1A). Differences remained in the distributions of mitochondrial and ribosomal RNA percentages, even after applying the respective thresholds (Supplementary Figures C-D), and no differences were observed between groups in terms of the number of genes or reads per cell (Supplementary Figures E-F). Nevertheless, cells with low number of genes detected (<300) and high number of molecules detected (>3,000) were excluded in order to avoid the inclusion of uninformative and/or doublet cells in the analyses, these thresholds were decided based on the distribution in Supplementary Figures 1E-F.

After applying all quality control filters, 82,698 cells remained. The gene profiling of these cells were normalized, scaled and the highest variable genes were selected using the Seurat R package^26^. To integrate the different samples into a common dimensional space, reciprocal principal component analysis (RPCA) was applied. Based on the RPCA results, a shared nearest neighbor (SNN) graph was constructed. Clustering was performed using the Louvain algorithm across multiple resolutions. To determine the optimal resolution, and thus the number of clusters, the stability of clusters across increasing resolutions was evaluated^27^. The resolution of 0.5, which yielded eleven stable clusters, was selected, as it provided the highest stability before cell assignments began to fluctuate significantly at higher resolutions.

### Single-nuclei RNA-Seq analysis of *S. senegalensis* male gonads identified the major cell types

Cell cluster annotation was performed based on manual inspection of cell type-specific markers (Supplementary Table 1) and pathway enrichment analysis of zebrafish MgSigDB^28^ selected gene sets (Supplementary Table 2). In the t-SNE representation (Figure 1A), all eleven clusters are clearly separated into the two-dimensional space. The heatmap of the top ten cell type markers (Figure 1B) shows distinct expression patterns for each cluster, supporting their specific identities.

**Figure 1:**
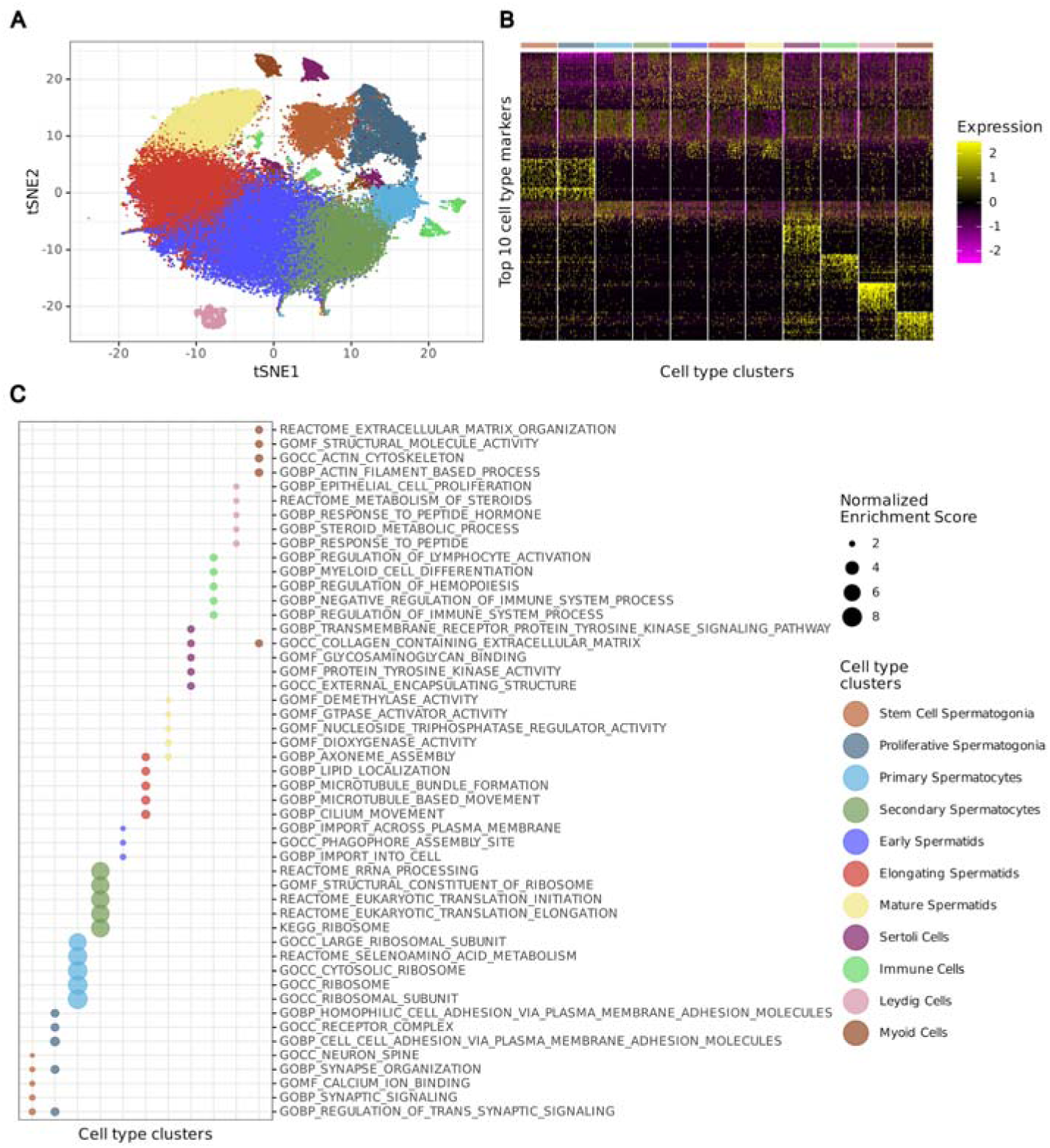
Cell type identification and functional characterization in scRNA-Seq of male gonad of Solea senegalensis. (A) t-SNE plot based on the top 50 principal components derived from the 2,000 most variable genes, cells are colored by identified cell type clusters. (B) Heatmap of the top 10 marker genes for each identified cell type cluster. Marker gene expression levels are scaled (Z-scores), with yellow indicating high expression and magenta indicating low expression. (C) Top 5 most enriched biological pathways for each cell type cluster (NES > 0 and q-value < 0.1). The size of each dot represents the normalized enrichment score (NES), while the color indicates the corresponding cell type. Pathways are sorted by cell types.

We identified two clusters with highly similar expression profiles (light brown and grayish blue clusters in Figure 1B). Both displayed elevated expression of genes associated with stem cell maintenance and self-renewal, such as the transcription factors *sox1a*, *grin2aa*, and *prdm16*, among others. Gene set enrichment analysis (GSEA) revealed that both clusters share enriched pathways related to synaptic signaling (Figure 1C). Although spermatogonia do not participate in synaptic signaling in the classical neuronal sense, elements of these pathways, such as neurotransmitters, receptors, and associated signaling proteins, may play a role in cell communication within the gonadal environment. Notably, the grayish blue cluster showed enrichment in functions related to cell-cell adhesion and receptor complexes (Figure 1C). Based on these observations, the light brown cluster likely represents spermatogonial stem cells, whereas the grayish blue cluster may comprise committed spermatogonia, which begin the differentiation process through strong interactions with Sertoli cells.

The five clusters with the largest number of cells are located at the center of the t-SNE plot (Figure 1A) and exhibit a continuous marker expression gradient (Figure 1B). The light blue cluster is characterized by high expression of ribosomal proteins, indicative of extensive protein synthesis, a hallmark of actively dividing cells. GSEA confirmed enrichment in ribosome-related processes and oxidative phosphorylation (Figure 1C), consistent with rapid cell proliferation and preparation for meiotic entry, features typical of primary spermatocytes. The dark green cluster also maintains high ribosomal protein expression along with translation factors, such as *eef1b2*, suggesting ongoing cell growth and readiness for meiosis. In this cluster, translational regulation and ribosome biogenesis are the most enriched pathways, along with pathways responsive to amino acid availability, hallmark of mid or secondary spermatocytes. The dark blue cluster appears to represent cells transitioning from late meiosis to early spermatids. It shows upregulation of genes related to post-transcriptional regulation (*cpeb1b*, *mbnl1*, *nova1*) and chromatin remodeling (*brdt*, *arid5a*, *kdm6ba*), indicating ongoing postmeiotic processes. Enrichment in pathways related to membrane trafficking and autophagy (Figure 1C) reflects the initiation of cellular remodeling, including uptake of essential metabolites and membrane components for flagellum and acrosome formation, while unnecessary organelles are being degraded. The red cluster is enriched in genes involved in cytoskeleton regulation and cell morphology (*myo10*, *arhgap21b* or *arhgap39*), as well as spermatid structural remodeling (*enah* or *spata17*). Enrichment in cilium movement, lipid location, and axoneme assembly pathways (Figure 1C) further supports its identification as elongating spermatids undergoing significant morphological changes. The yellow cluster continues to express structural remodelling but also shows markers associated with sperm maturation and acrosome development (*dcst1*, *cadm3* or *tspan4a*). Functional enrichment in GTPase cycle and phosphatase activity suggest these cells are undergoing terminal remodelling, typical of late spermatids completing spermiogenesis (Figure 1C).

The remaining clusters correspond to non-germline male gonad cell types. Sertoli cells (purple cluster) are identified by enrichment in structural, signalling and adhesion-related pathways (Figure 1C), including the key marker *nr5a2*, which encodes a nuclear receptor essential for Sertoli cell function. Immune cells (light green cluster) are characterized by immune-related pathways associated with both lymphoid and myeloid lineages (Figure 1C). Specific markers include *ptprc* (a pan-leukocyte antigen), *cd74b* (a B-cell marker), and *cd68* (a macrophage marker), among others. Leydig cells (pink cluster) are identified by pathways involved in steroid biosynthesis and hormone production (Figure 1C). Marker genes such as *star* (steroidogenic acute regulatory protein), *ar* (androgen receptor), and *hsd11b2* (involved in steroid metabolism) confirm this identity. The brown cluster likely represents peritubular myoid cells or other structural cells contributing to the extracellular matrix (ECM) maintenance and testicular integrity. The expression of *tpm1*, *col6a3*, and *cald1b*, genes involved in smooth muscle function and ECM composition, supports this allocation. Enriched pathways also align with structural roles in supporting seminiferous tubules and facilitating sperm transport (Figure 1C).

### Trajectory analysis based on single-nuclei RNA-Seq molecular data revealed a spermatogenesis differentiation trajectory from marker-annotated spermatogonial stem cell to mature spermatids

Germline cell annotation was not an easy task based on functional markers, first due to the lack of functional annotation for many *S. senegalensis* genes that appears as strong markers of many of the clusters but without known function, and second because the spermatogenesis is a continuous process where frontiers between cell types are hard to define and thus some markers may overlap similar cell types. Therefore, in order to verify the existence of a transcriptional continuum between stem cell spermatogonial cells and the mature spermatids, germline cells were selected from the previous analyses and a trajectory inference was performed using Monocle3^29^.

The analysis revealed one single significant trajectory, without branches, flowing from the stem cell spermatogonial cluster to the mature spermatids cluster (Figure 2A), as defined by the functional marker-based annotation in the previous analysis. To define the sequence of regulatory molecular changes driving the cell differentiation along spermatogenesis, four principal coregulated gene modules were identified (Figure 2B). The functionality of these four modules were tested based on gene overrepresentation analysis on MgSigDB selected gene sets. Three of these modules (1, 2 and 4 in Figure 2C) showed significant overrepresented pathways (q-value < 0.1 and enrichment score > 0). Module 4 was overexpressed in spermatogonial cells with a peak at the proliferating spermatogonia; then its expression decreased after primary spermatocytes (Figure 2B), showing pathways related to ribosomal biogenesis and rRNA maturation, consistent with increased translational demand during early differentiation. Additionally, responsiveness to fibroblast growth factors and regulation of ubiquitin-mediated proteolysis highlight active engagement with proliferative and survival signals, positioning these cells for mitotic expansion and lineage commitment (Figure 2C). Module 2 peak corresponds to spermatocytes; the expression of cluster-specific markers increases since proliferative spermatogonia and decreases at early stages of spermatid maturation (Figure 2B). Unexpected enrichment of steroid hormone biosynthesis and glucocorticoid metabolism pathways in spermatocytes suggests that these cells may transiently engage in local steroidogenic activity or modulate their sensitivity to hormonal process. Additionally, enrichment of potassium channel regulation indicates active ionic remodeling during meiotic progression, consistent with known changes in membrane dynamics and volume regulation (Figure 2C). Module 1 defines the last stages of the differentiation, the expression initiates at secondary spermatocytes, reaching its peak at early spermatids but maintaining a high expression in all spermatid clusters (Figure 2B). In this module, pathways related to oxidative phosphorylation and mitochondrial ATP synthesis were enriched, consistent with the increased energy demands associated with sperm motility. Additionally, omega peptidase activity suggests active proteolytic remodeling during the final stages of spermiogenesis (Figure 2C).

**Figure 2:**
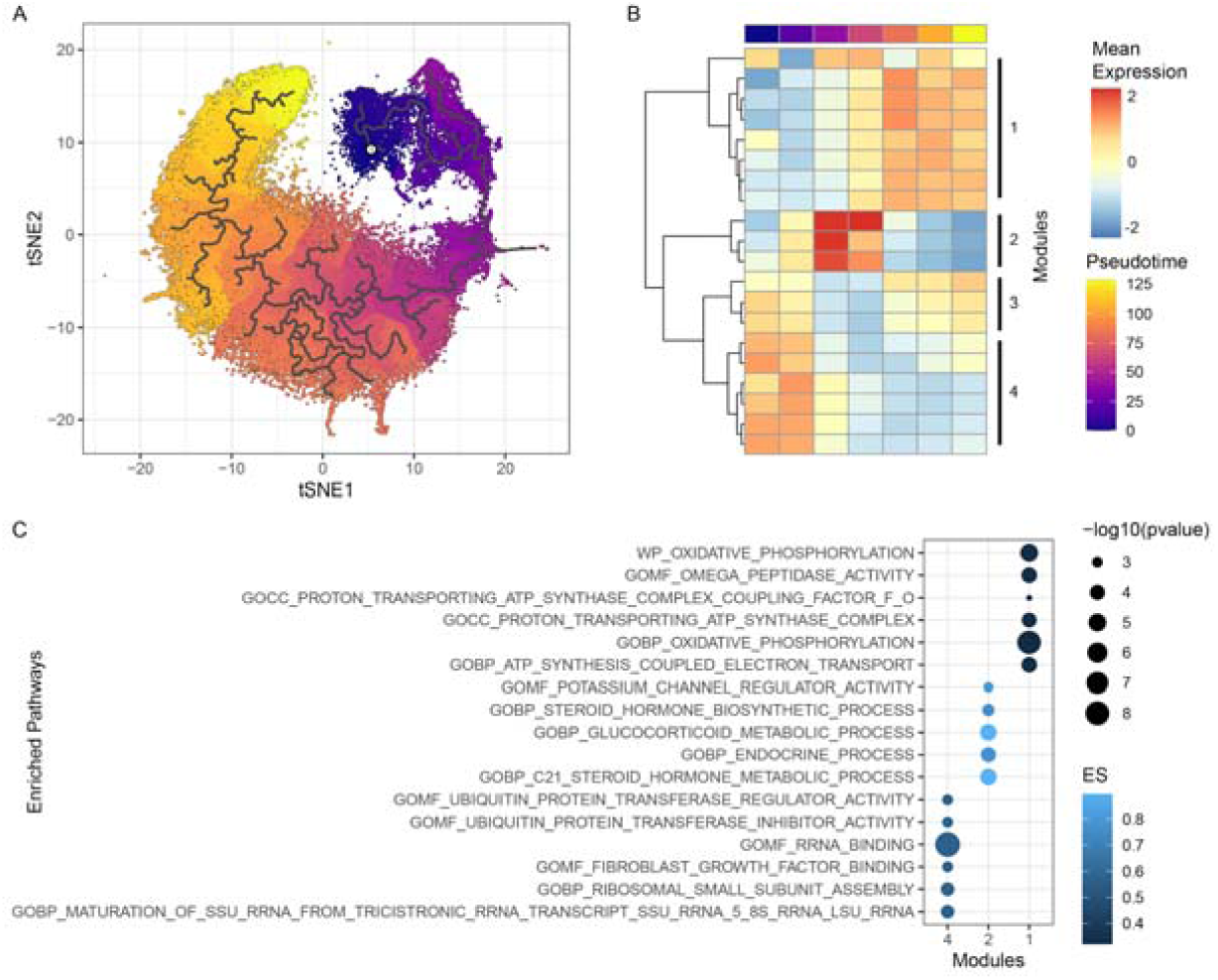
Pseudotime trajectory and module-specific gene expression dynamics on germline male gonad cells. (A) t-SNE plot showing the distribution of cells along pseudotime, colored by pseudotime progression from purple (early) to yellow (late). Black lines represent the inferred trajectory path across the cell population. (B) Heatmap showing the expression patterns of genes grouped into co-expression modules across pseudotime. Rows correspond to gene modules, and columns represent each of the identified cell types. Gene expression is averaged and scaled, and the color bar above the heatmap indicates average pseudotime progression per cell type. (C) Top 5 most overrepresented pathways for the genes contained in each gene module identified during trajectory analysis (ES > 0 and q-value < 0.1). The size of the dots represents the statistical significance of enrichment (– log10(p-value)), while the color intensity denotes the enrichment score (ES), with lighter colors indicating stronger enrichment.

### Captive-bred males showed overrepresentation of spermatogonial cell stages compared to wild-bred males

There are different types of infertility, and some may be associated with alterations in the cellular composition along spermatogenesis, such as in cases of azoospermia or oligospermia. To investigate cell type composition through differentiation, we visualized overall cell type density using t-SNE dimensionality reduction (Figure 3A) and represented individual cell type proportions using bar plots (Figure 3B). Both analyses revealed overrepresentation of spermatogonial cells, particularly proliferative spermatogonia, and a depletion of mature spermatids in CB males. To validate the observed imbalance in cell proportions, we performed bulk RNA-Seq analysis on independent biological samples. GSEA was conducted using bulk RNA-Seq differential analysis results and the top 100 marker genes for each germ cell type involved in spermatogenesis. CB animals showed a significant depletion of spermatid markers (NES < −2), with p-values < 5e-10 across all three identified spermatid subtypes, as well as an enrichment in spermatogonial markers (NES ∼ 1), but not statistically significant (Supplementary Table 4). Notably, a clear transition from marker enrichment to depletion across differentiation stages was evident when plotting the running enrichment score for all cell types (Figure 3C).

**Figure 3.**
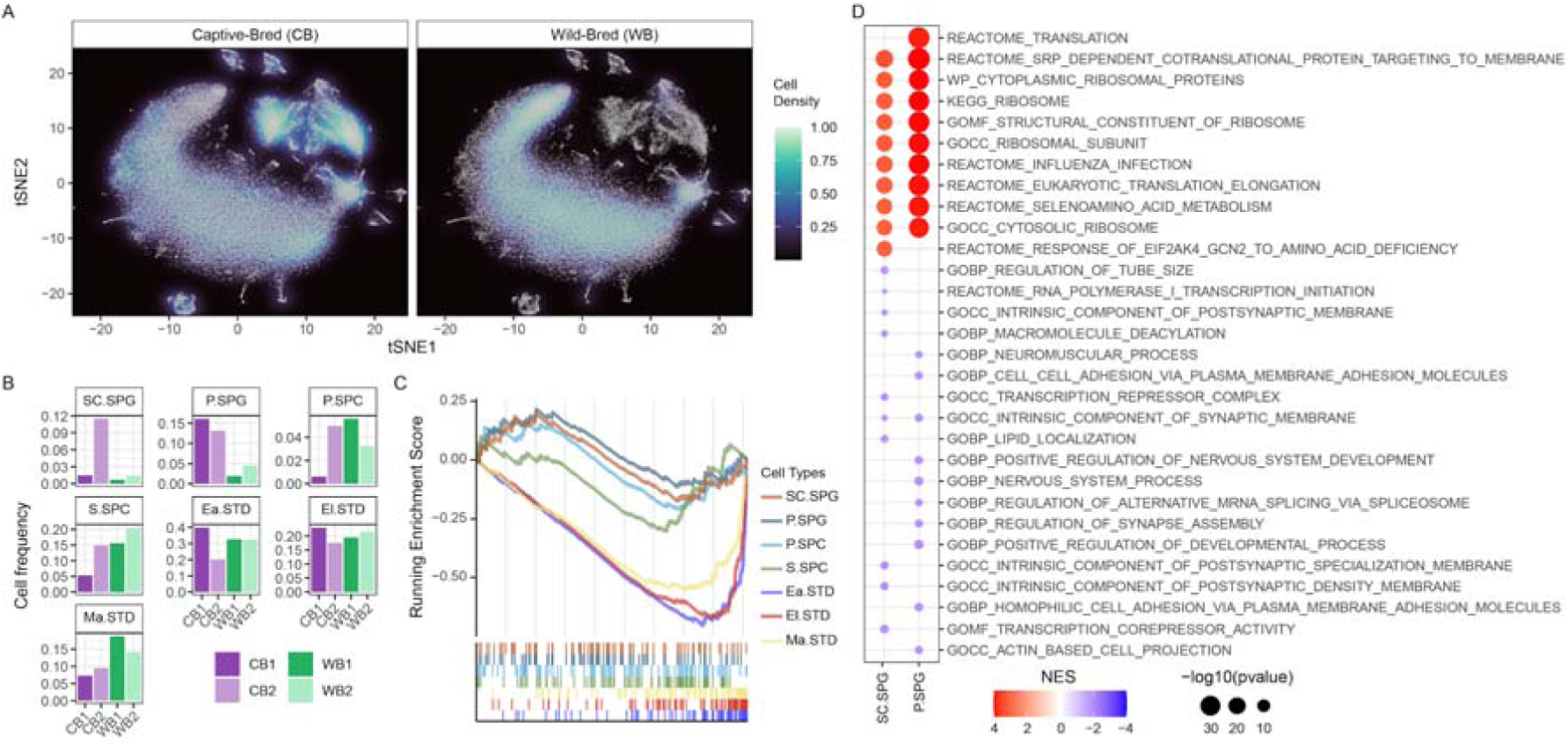
Altered cell type composition and cell-cell interactions in gonads from CB males. (A) t-SNE projections of scRNA-Seq from male gonads of CB and WB individuals, showing cell density. (B) Bar plots representing the relative frequency of each annotated germ cell type across biological replicates for both CB and WB groups. Cell types are SC.SPG (stem cell spermatogonia), P.SPG (proliferative spermatogonia), P.SPC (primary spermatocytes), S.SPC (secondary spermatocytes), Ea.SPD (early spermatids), El.SPD (elongating spermatids), Ma.SPD (mature spermatids). (C) GSEA results based on differential analysis of male gonad bulk RNA-Seq data between CB and WB individuals. Enrichment scores were calculated using the top 100 marker genes for each germ cell identified in the scRNA-Seq. (D) Dot plot showing the top five upregulated and downregulated pathways from the GSEA analysis comparing CB and WB groups in stem cell spermatogonia (SCSg) and proliferative spermatogonia (PSg). Dots are colored by normalized enrichment score (NES) and sized by −log10(q-value).

The overrepresentation of spermatogonial cell types in CB animals suggests the existence of molecular dysregulation that impairs the commitment of spermatogonial cells to the spermatogenic differentiation process. To investigate which molecular pathways may be involved in this impairment, differential expression analysis and GSEA were performed on the stem cell and proliferative spermatogonial clusters, comparing CB and WB animals. The analysis revealed two major groups of downregulated pathways in CB spermatogonial cells: those related to synaptic and neural processes, and those associated with adhesion and cytoskeletal dynamics (Figure 3D and Supplementary Table 5). As previously mentioned, although synaptic and neuronal terms are classically associated with the nervous system, similar mechanisms are active in spermatogonial cells, particularly in cell communication and differentiation. Their downregulation may therefore indicate impaired cell communication, potentially disrupting the differentiation process. Additionally, the downregulation of adhesion and cytoskeleton-related pathways may compromise the interaction between spermatogonial and Sertoli cells, which is essential for niche maintenance and proper differentiation. In contrast, the upregulated pathways in CB spermatogonial cells were mainly associated with ribosome structure, translation, and co-translational protein targeting, suggesting a shift in cellular activity toward increased protein synthesis.

### Spermatogonia-Sertoli cell interactions are markedly dysregulated in captive-bred males

To further investigate the potential role of downregulated cell-cell communication during spermatogonial stages, NicheNet^30^ was employed to assess intercellular signaling in proliferative spermatogonia. Since the NicheNet cell-cell communication network is based on human and mouse data, only orthologous genes between *S. senegalensis* and humans, as identified in the NCBI orthologs database^31^, were included in the analysis. Briefly, NicheNet calculates the prior interaction potential of ligand-receptor pairs using biological knowledge gathered from multiple databases. It then estimates ligand activity based on how well the expression of a ligand in ligand-producing cells (senders) can explain observed expression changes in target genes within the receiver cell population. Finally, it computes a regulatory potential score, representing the likelihood that a ligand influences a downstream target gene through intermediate signaling and transcriptional networks.

A differential expression analysis of target genes in proliferative spermatogonia from the NicheNet database revealed a substantial downregulation in CB animals (Fig. 4A), with over 50% of target genes being downregulated. This observation is consistent with our previous findings linking gene downregulation to impaired cell-cell communication. These downregulated target genes were subsequently used for ligand activity calculations. Sertoli, Leydig, myoid, and immune cells were considered potential senders interacting with spermatogonial receptors. Accordingly, the database was filtered based on the expression of ligands in senders and receptors in the receiver cell. Ligand-receptor pairs were then selected based on this filtering, and their prior interaction potential was estimated. Among these, the receptor with the highest number of potential interactions affecting downregulated target genes was the epidermal growth factor receptor (*egfr*) (Fig. 4B), a gene known to play a crucial role in spermatogonial stem cell differentiation.

**Figure 4.**
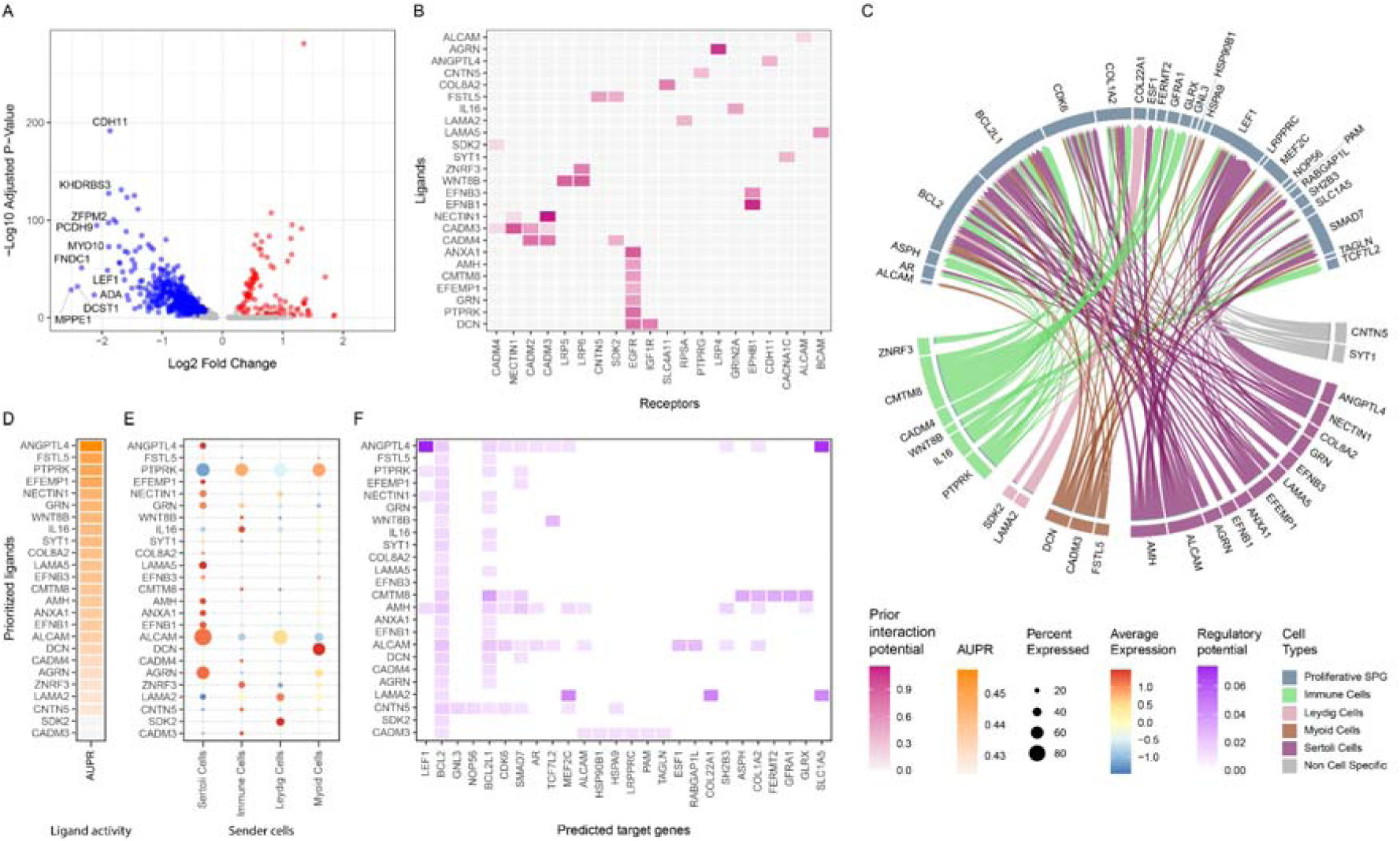
Cell-cell communication networks reveal downregulated target genes downstream of ligand–receptor interactions. (A) Volcano plot showing differentially targeted genes downstream of ligand–receptor interactions between CB and WB animals. Significantly upregulated genes are shown in red, and downregulated genes in blue. (B) Heatmap of ligand-receptor pairs upstream of downregulated target genes in CB animals, highlighting prior interaction potential. (C) Circos plot displaying inferred ligand-target networks. Links represent prioritized signaling interactions from sender to receiver cells, colored by the predominant ligand-expressing sender cell type. P.SPG stands for proliferative spermatogonia. (D) Heatmap ranking ligands by their activity scores (AUPR), based on the downregulated target genes. (E) Dot plot showing ligand expression across sender cell types. Dot size represents the percentage of cells expressing each ligand, while color denotes average expression level. (F) Heatmap showing the regulatory potential of ligands on predicted target genes.

Then, ligand-producing cell types were identified by comparing the expression levels of each ligand across all potential senders. A ligand was considered specifically produced by one sender if it showed a mean expression level higher than the mean plus one standard deviation across all potential senders. This analysis showed that nearly half of the ligands (12 out of 25) were predominantly expressed in Sertoli cells. Consistently, nearly half of the target genes affected by these interactions with spermatogonial cells receptors (75 out of 152) were also mediated by these Sertoli cell-derived ligands (Fig. 4C). The target genes with the highest number of regulatory interactions originating from Sertoli cells were *bcl2*, *bcl2l1*, and *cdk6*, each with eleven interactions, followed by *lef1* and *smad7*, each with ten. Additionally, although involved in fewer interactions, key genes involved in spermatogenesis, such as *ar* (androgen receptor), were also affected by ligand-receptor dysregulation (Fig. 4C). Once ligand activity was inferred based on target gene downregulated in the proliferative spermatogonia of CB animals, *angptl4* emerged as the ligand with the highest activity (Fig. 4D). It is almost exclusively expressed in Sertoli cells (Fig. 4E) and exhibits a strong regulatory potential over many of the genes that are also highly targeted by other Sertoli cell-produced ligands (Fig. 4F). These include *lef1*, the target gene with the highest regulatory potential; *bcl2*, the gene influenced by the greatest number of ligands with positive regulatory potential; as well as other key genes such as *ar* and *slc1a5*, the latter being the second highest in regulatory potential among the target genes.

## Methods

### Biological sample acquisition

Adult CB and WB males were sacrificed by decapitation at the SEA EIGTH farm (Safiestela, Portugal) following the ethical procedures of the company. Both gonads from each animal were dissected and immediately flash-frozen in liquid nitrogen, to be then stored at −80°C until their use for the single-nuclei protocol. One testicle from each individual was used for snRNA-Seq and the other for bulk RNA-Seq. In total, for snRNA-Seq 2 CB and 2 WB animals were available, while for bulk RNA-Seq the analysis was performed on 3 CB and 6 WB animals.

### Molecular profiling

#### Single-nuclei RNA-Seq library preparation

Nuclei were extracted following a published protocol^32^ with small modifications for use in non-standard vertebrate species^33^. Briefly, the frozen gonads were homogenized using a micropestle in 400 µL ice-cold homogenisation buffer (250mM sucrose, 25mM KCl, 5mM MgCl2, 10 mM Tris-HCl [pH 8], 0.1% IGEPAL, 1 µM dithiothreitol [DTT], 0.4 U/µL Murine RNase Inhibitor [New England BioLabs, M0314L], and 0.2 U/µL SUPERase-In [Ambion, AM2694]). The homogenates were triturated gently using a p1000 tip 10 times, incubated on ice for 5 min and then centrifuged at 100 g for 1 min at 4 °C to pellet any unlysed tissue chunks. The supernatant was transferred into another 1.5mL Eppendorf tube and centrifuged at 400 g for 4 min at 4 °C to collect nuclei. The nuclei were washed twice in 400 µL homogenization buffer and strained using a 40 µm Flowmi strainer (Sigma, BAH136800040) during the second wash step to remove nuclei aggregates. The final nuclei pellet was resuspended in 30–50 µL Nuclei Bufer (10x Genomics, PN-2000207). To estimate the nuclei concentration, nuclei aliquots were diluted in phosphate-buffered-saline (PBS) with Hoechst and propidium iodide (PI) DNA dyes and counted on Countless II FL Automated Cell Counter (Thermo Fisher Scientific, RRID: SCR_020236). The Chromium Next GEM Single Cell 3’ Reagent Kits v3.1 (PN-1000121, PN-1000120, and PN-1000213) were used to make snRNA-seq libraries. Libraries were quantified on a Qubit Fluorometer (Termo Fisher Scientifc; RRID: SCR_018095) and quality checked on a Fragment Analyzer (Agilent; RRID: SCR_019417). Finally, single-nuclei RNA-seq libraries were sequenced on an Illumina NovaSeq 6000 platform, generating 150 bp paired-end reads.

#### Bulk RNA-Seq library preparation

After sample thawing, total RNA was extracted and purified using the miRNeasy Kit (QIAGEN). RNA integrity and quantity were evaluated in a Bioanalyzer (Bonsai Technologies, Madrid, Spain) and in a NanoDrop® ND-1000 spectrophotometer (NanoDrop® Technologies Inc., Wilmington, DE, USA). RNA samples were delivered to Novogene (UK) for library preparation using NEBNext Ultra Directional RNA Library Prep Kits for Illumina and sequenced using an Illumina NovaSeq S4 platform to generate 150 bp paired end reads.

### Statistical analysis

#### Single-nuclei RNA-Seq quality control and processing

The *S. senegalensis* IFAPA_SoseM_1 genome assembly (GCF_019176455.1) and corresponding gene annotation were downloaded from the NCBI RefSeq Genome Database^34^. A CellRanger (v8.0)^35^ reference was constructed, and FASTQ files were subsequently processed, aligned, and annotated. Additionally, CellRanger filtered cell barcodes to remove empty droplets, cellular debris, and ambient RNA, retaining barcodes that represented true cells.

For downstream analysis, cells were retained if they exhibited more than 500 counts, and genes were retained if they had more than 3 counts in at least 50 cells. Additional filtering criteria based on mitochondrial and ribosomal gene percentages, as well as total counts and feature counts per cell, were applied as follows: mitochondrial gene percentage <10%, ribosomal gene percentage >1%, gene count per cell >300, and total counts per cell <3,000.

Cell cycle scores were calculated using the Seurat R package (v5.1)^36^, employing predefined reference gene sets^37^. Gene expression data were normalized and scaled using Seurat, and the top 2,000 most variable features were identified using the variance-stabilizing transformation ("vst") method. Dimensionality reduction was performed using principal component analysis (PCA), retaining the first 50 principal components (PCs). Integration of scRNA-seq samples was carried out using reciprocal PCA (RPCA). Subsequently, t-distributed stochastic neighbor embedding (t-SNE) was applied to reduce the first 50 RPCA dimensions to two-dimensional space for visualization.

To identify the most stable clustering configuration, a k-nearest neighbor (k-NN) graph was constructed based on the first 50 RPCA dimensions to capture relationships between cells. Community detection using the Louvain algorithm was performed at multiple resolutions (0.05, 0.1–1.0 in increments of 0.1, 1.5, and 2.0). The resulting Louvain clustering resolutions were compared using the clustree R package^38^ to determine the optimal, most stable cluster configuration.

#### Bulk RNA-Seq quality control and processing

Briefly, raw reads quality was examined using FASTQC^39^ and adapters were trimmed using Trim Galore^40^, a wrapper tool of Cutadapt^41^. Filtered reads were then mapped to the *S. senegalensis* genome using STAR^42^. For differential expression analysis, normalized transcript reads were filtered based on their expression, preserving only genes with a Transcript per Million Reads (TPM) value > 5.

#### Functional and enrichment analyses

Functional interpretation of marker genes and differential expression analyses were conducted using Gene Set Enrichment Analysis (GSEA) and Overrepresentation Analysis (ORA) implemented in the ClusterProfiler R package (v4.12.6)^43^. Orthologs of Solea genes in Zebrafish were identified via NCBI Gene IDs and the NCBI ortholog database^44^ using the Orthology.eg.db R package (v3.19.1). Zebrafish gene sets from the Molecular Signatures Database (MsigDB)^45^, retrieved using the msigdbr R package (v7.5.1), were utilized for functional analyses. The gene sets included BIOCARTA, KEGG, PID, Reactome, WikiPathways, and Gene Ontology (GO) terms.

#### Single-nuclei trajectory analysis

Single-nuclei trajectory analysis was performed using Monocle3^29^, following the authors’ guidelines. Cell clusters corresponding to stages within the spermatogenesis differentiation process were selected, and t-SNE RPCA embeddings from the complete dataset were utilized for this analysis. The trajectory root was programmatically defined, as recommended by the authors, within the spermatogonial stem cell cluster. To identify correlated gene modules, genes exhibiting variation across the trajectory were selected based on Moran’s I spatial autocorrelation metric (q-value < 1e-5). Louvain community detection was then applied to gene embeddings at multiple resolutions (1e-6, 1e-5, 1e-4, 1e-3, 1e-2, 1e-1) to determine the most stable partition, resulting in the identification of twenty co-regulated gene modules. These twenty modules were further grouped into four principal modules based on hierarchical clustering of average expression across modules and cell types identified in the analysis.

#### Differential and cell-cell interaction analyses

Bulk RNA-Seq differential analysis was conducted using the R package DESeq2 (v1.38.1)^46^ comparing CB versus WB animals.

Single-nuclei RNA-seq differential expression analyses were performed using the Wilcoxon test implemented in Seurat. Cluster marker genes were identified by comparing cells within a cluster against all other cells, while functional differential analyses compared cells of the same cell type between different conditions (CB versus WB).

Cell-cell interaction analyses were performed using the NicheNet R package (v2.2.0)^30^. Given that the cell-cell interaction database curated by the NicheNet authors only contains human and mouse datasets, this analysis was restricted to Solea genes with human orthologs identified via NCBI Gene IDs and the NCBI ortholog database^44^ using the Orthology.eg.db R package (v3.19.1).

## Discussion

Selective breeding programs in aquaculture have proven highly effective across numerous orders of finfish and shellfish, enabling producers to meet consumer demand competitively while maintaining ecological integrity and preserving biodiversity^47^. Flatfish have been successfully bred in captivity worldwide, achieving significant economic outcomes. However, for *S. senegalensis*, one of the most promising European flatfish species due to its flesh quality and high market value^10^, genetic breeding programs aimed at enhancing productivity face important challenges related to reproductive constraints on farms. This has primarily to do with CB males, which produce low and variable sperm volumes^12–14^. Furthermore, these males are unable to perform the typical courtship with females^10,11^, which suggests a connection with their low sperm production. Neither CB females nor WB males maintained under captivity conditions exhibit reproductive limitations^15,48^, which prompts to focus the problem on CB males.

Previous studies employed gonadal bulk RNA-Seq and methylation profiling, they revealed numerous differences between CB and WB males and identified critical pathways associated with male gonadal development, including MAPK signaling, Wnt signaling pathway, olfactory receptors, focal adhesion, or ECM-receptor interactions, among others^49–51^. However, bulk molecular analyses in complex biological samples containing diverse cell types and variable cell numbers, without an effective deconvolution signature matrix, provide limited information regarding the specific cell types in which dysregulation occurs and fail to capture the complete molecular landscape underlying infertility^52^. Thus, to elucidate the molecular mechanisms underlying infertility in captive-bred males at the cellular level, gonadal tissue from adult CB and WB males was collected and subjected to snRNA-Seq profiling.

Single-nuclei RNA sequencing enabled the identification of eleven distinct cell clusters within the gonads of *S. senegalensis*. These clusters encompassed multiple stages of the spermatogenesis process, including spermatogonial stem cells, differentiating spermatogonia, spermatocytes, and spermatids. Additionally, we identified supporting cell types such as Sertoli cells, endocrine cell populations including Leydig cells, and, at lower abundances, structural and immune cell populations within the gonadal environment^53,54^. All identified cell clusters exhibited the expression of well-established marker genes, confirming their cellular identities within the dataset. However, a substantial number of genes demonstrated high cluster specificity beyond known markers, suggesting their potential as candidate genes for further functional characterization. These cluster-specific genes could serve as valuable tools for developing accurate signature matrices for future deconvolution analyses in bulk transcriptomics, aiding the monitoring of gonadal status and reproductive health in *S. senegalensis* and related species.

Trajectory analysis of spermatogenesis further elucidated the key dynamic molecular pathways underlying the differentiation process from spermatogonial stem cells to mature spermatids. We observed that pathways associated with ribosome biogenesis, growth factor signaling, and ubiquitination processes were highly active during the spermatogonial and early spermatocyte stages, consistent with the high proliferative and translational demands during the initial phases of germ cell development. These processes declined as cells transitioned into later spermatocyte stages, where we observed a marked enrichment in endocrine and hormone biosynthetic pathways, reflecting the critical role of steroid hormones in regulating meiosis and germ cell maturation^55^. In the spermatid clusters, pathways associated with oxidative phosphorylation and ATP synthesis were prominently upregulated, aligning with the increased energy demands required for chromatin condensation, flagellar development, and cellular remodeling during the final stages of spermiogenesis^56^. Collectively, these findings highlight the dynamic and stage-specific molecular landscape of spermatogenesis in *S. senegalensis*, providing a foundational framework for understanding the molecular basis of male fertility in this species.

Among the potential causes of male infertility, some can be identified by examining the distribution of cells across the different stages of spermatogenesis. In the case of CB *S. senegalensis* males, we observed the presence of all spermatogenic cell types. However, their proportions differed markedly from those in WB counterparts. This indicates that male infertility in CB individuals is not due to a complete arrest of the spermatogenesis process but rather to a limitation in the number of cells successfully completing differentiation. Specifically, undifferentiated cell types, such as spermatogonial stem cells and proliferative spermatogonia, were overrepresented in CB males, while mature spermatids were 2-3 times less abundant compared to WB males. This skewed distribution suggests that the molecular disruptions underlying impaired reproductive success in captivity result in oligospermia, characterized by reduced numbers of mature sperm cells despite the continuation of spermatogenesis. It is noteworthy that while in vitro fertilization using sperm from CB males is technically feasible, the low sperm yield makes it economically impractical for implementation in large-scale breeding programs^12–14^.

To elucidate the molecular and endocrine mechanisms constraining the completion of spermatogenesis in captivity, we performed a differential analysis of spermatogonial clusters between CB and WB *Solea senegalensis* males. The results revealed a marked downregulation of molecular pathways related to cell-cell communication and cell adhesion processes in CB males, consistent with previous findings at the bulk RNA-Seq level^51,57^. Due to limited annotation of fish-specific genes involved in cell-cell communication and the absence of experimentally validated ligand-receptor databases in teleosts, our analysis relied on known human orthologs to predict potential ligand-receptor interactions. Strikingly, more than 50% of the predicted downregulated signaling pathways specifically involved Sertoli cell to proliferative spermatogonial cell interactions, indicating a significant impairment in this communication, which is crucial for the progression of spermatogonial differentiation. Sertoli cells are the only somatic cells in direct contact with spermatogonial cells, and proper cell-cell communication is essential for testis formation and spermatogenesis^58^.

Among the ligand-receptor pairs implicated, several ligands known to be highly expressed in Sertoli cells, including *nectin1*, *efnb1*, *efnb3*, *agrn* and *lama5*, were identified. These ligands are well-established mediators of Sertoli-spermatogonial adhesion and signaling. Additionally, Anti-Müllerian Hormone gene (*amh*), critical for male gonad development and Sertoli cell maturation, was also identified as an affected ligand in our cell-cell communication analysis. On the receiving end of these impaired interactions, genes associated with the self-renewal and maintenance of spermatogonial stem cells, such as *gfra1*, as well as *lef1* and *tcf7l2* (Wnt signaling mediators), were notably impacted. Moreover, ar (androgen receptor), a key regulator of spermatogenesis progression, was also identified among the affected downstream targets. Interestingly, while some identified genes have not been directly associated with spermatogenesis, they may still play important roles in this context. Notably, *angplt4* exhibited the highest predicted ligand activity in our analysis, suggesting a potential involvement in Sertoli-germ cell communication and gonadal function that warrants further investigation.

In addition, *bcl2* and *bcl2l1* emerged as the two genes predicted to be regulated by the highest number of interactions. These genes are known to regulate spermatogonial survival and apoptosis under the influence of Sertoli-derived paracrine factors^59^. The impaired regulation of bcl2 and bcl2l1 is particularly significant, as defective apoptotic processes in spermatogonial cells unable to progress through spermatogenesis may lead to their accumulation. This accumulation could, in turn, further disrupt Sertoli-spermatogonial interactions, creating a self-reinforcing loop that blocks progression to the subsequent stages of differentiation, thereby contributing to the oligospermia observed in CB males.

Further investigations will be required to elucidate how these Sertoli-spermatogonial interactions become initially impaired, leading to the accumulation of undifferentiated spermatogonial cells in CB males. Notably, the oligospermia observed in these animals does not occur during the acclimation of WB males to captive conditions^11,15^. This observation leads us to hypothesize that epigenetic processes occurring during *S. senegalensis* development under captive conditions may result in defective gonadal maturation of CB males. Despite the limited number of individuals profiled in this study and the absence of species-specific ligand-receptor interaction databases, our single-nuclei transcriptomic analysis was able to identify defective Sertoli-spermatogonial interactions as a key molecular driver of oligospermia in CB *S. senegalensis* males. These impaired interactions involve critical genes associated with spermatogonial differentiation and apoptosis regulation, leading to the accumulation of undifferentiated, defective spermatogonial cells within the gonads. Our findings provide a clear molecular basis for understanding the reproductive dysfunction observed in CB males, offering a framework for future studies to target the molecular and epigenetic mechanisms constraining the successful completion of spermatogenesis in captivity.

## Acknowledgements

This study was funded by Proyectos Plan Operativo FEDER Andalucía 2021-2027 (C-EXP-159-UGR23), MICIU/AEI/10.13039/501100011033 project (PID2022-137821OB-C31) and by Consellería de Economía, Industria e Innovación e Consellería de Cultura, Educación, Formación Profesional e Universidades, Xunta de Galicia (ED431C 2022/33). DTS was supported by a regional research fellowship (Xunta de Galicia, 06_IN606D_2022_2693134), MC by a fellowship within the framework of the State Plan for Scientific, Technical and Innovation Research 2021-2023 (PID2022-137821OB-C31) and GB by MICINN, Juan de la Cierva-Incorporación (IJC2020-043364-I).

## Data Availability Statement

Single-nucleus RNA-Seq FASTQ and H5DF files will be available in the GEO repository. Bulk RNA-Seq data are currently being prepared for a separate publication and are available upon request. These data will be made publicly available following publication.

## Supplementary Figures

**Supplementary Figure 1:**
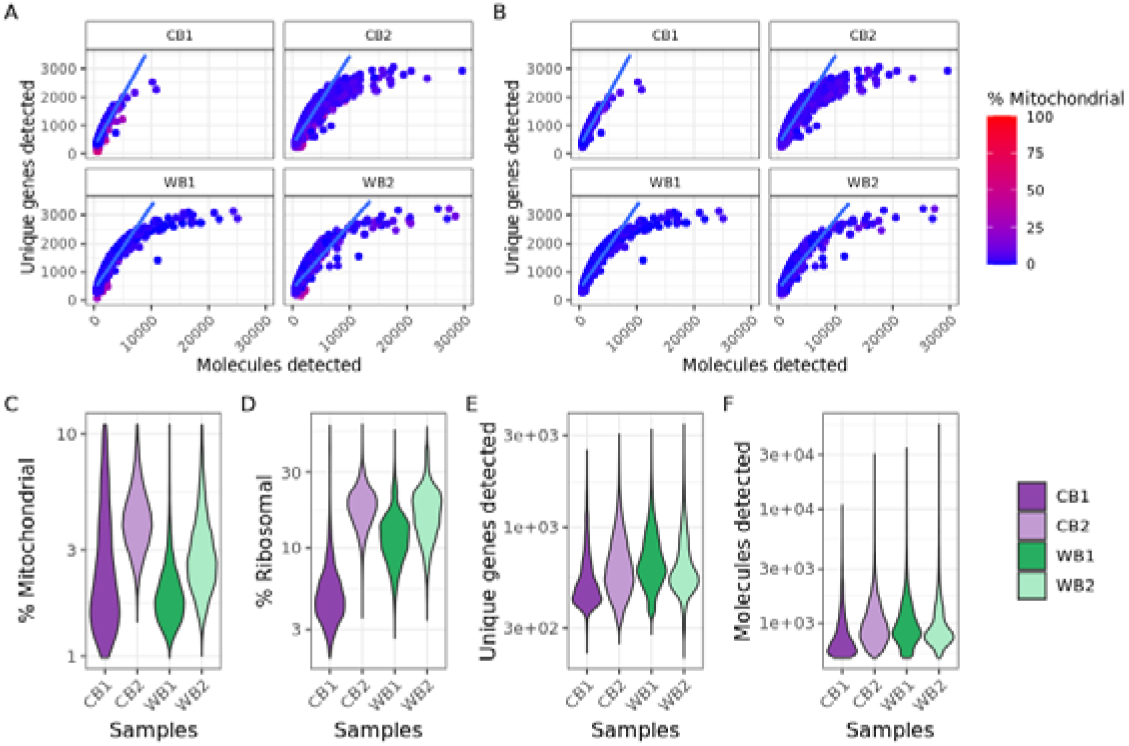
The number of molecules versus the number of unique genes (covered by at least one read) is plotted per cell (A) before and (B) after applying mitochondrial and ribosomal percentage thresholds (% mitochondrial < 10 and % ribosomal > 1). Each cell is colored based on its mitochondrial percentage, ranging from blue to red, with a midpoint at the 10% threshold. And the best fit line for a linear regression model between the number of unique genes and molecules detected is depicted in blue. Distributions of key quality metrics after thresholding are shown: (A) mitochondrial and (B) ribosomal percentages, (C) number of unique genes detected, and (D) number of molecules detected. CB and WB distributions for each sequenced individual are represented in purple and green tones, respectively.

**Supplementary Figure 2:**
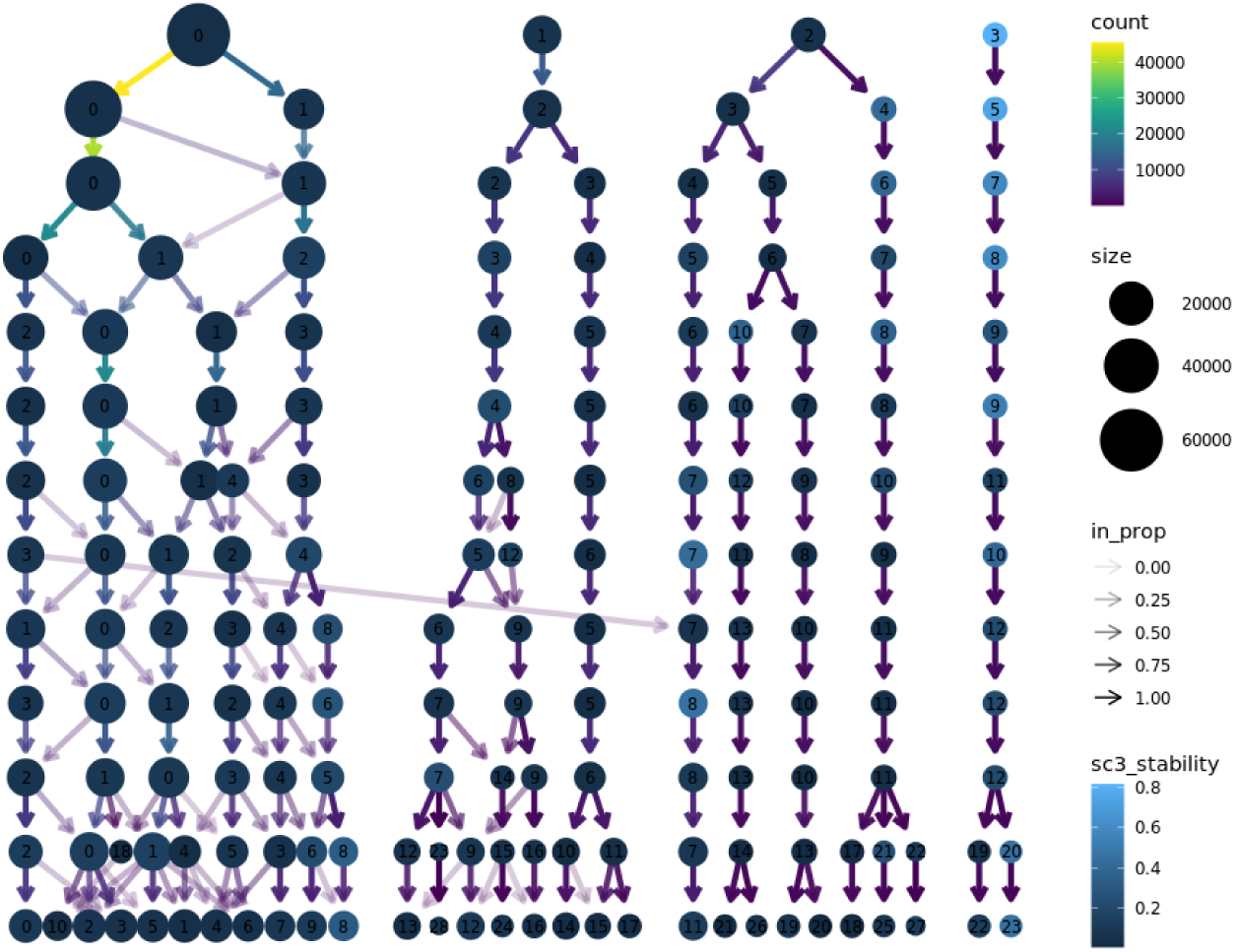
A clustering tree for shared nearest neighbor clustering algorithm results for resolutions 0.05, 0.1 to 1 with 0.1 step, 1.5 and 2 is shown. Each row of dots represents the clusters for each resolution; the size of the dots indicates the number of cells grouped within each cluster, and the blue color scale reflects the stability of each cluster across resolutions. The color of the arrows represents the number of cells shared between clusters at different resolutions, while their transparency indicates the proportion of shared cells.

**Supplementary Figure 3:**
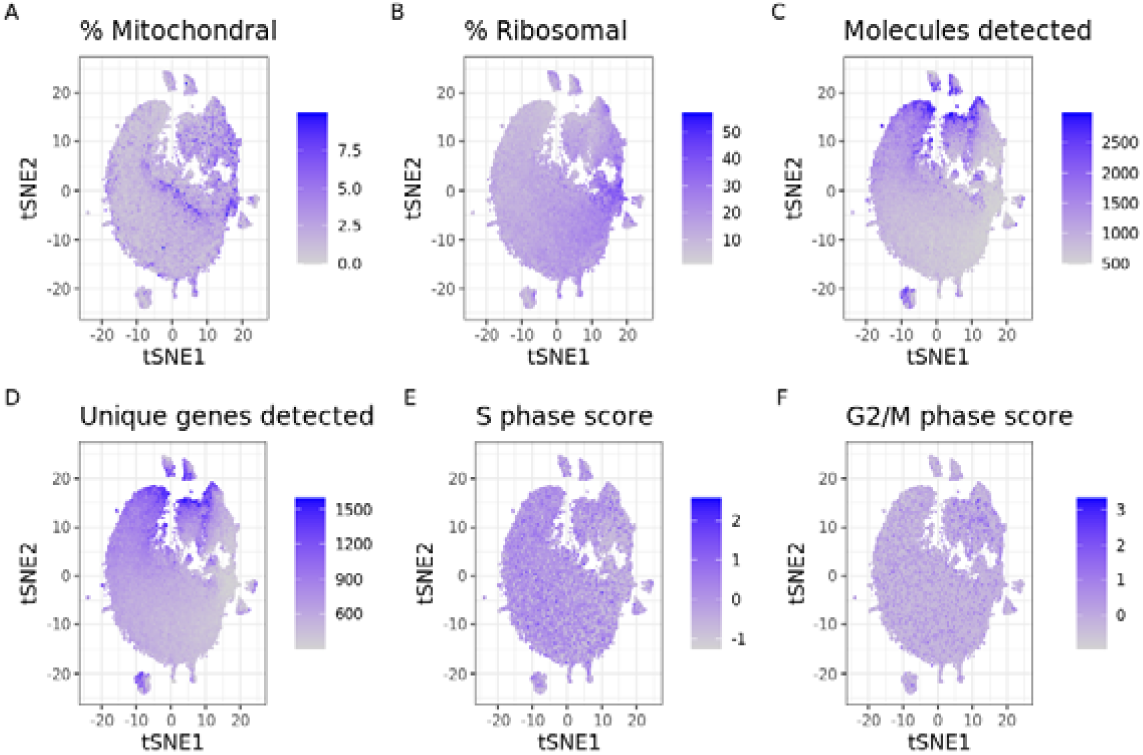
tSNE dimensional reduction based on the top 50 principal components derived from the 2,000 most variable genes, cells are colored based on quality metrics: (A) mitochondrial percentage, (B) ribosomal percentage, (C) number of molecules detected, (D) number of unique genes detected, (E) S phase score estimation and (F) G2/M phase score estimation.

